# Soil texture regulates bacterial motility and chemotactic recruitment to plant roots

**DOI:** 10.1101/2025.09.08.674882

**Authors:** Ahmed Al Harraq, Gayoung Choi, Sujit S. Datta, Joshua W. Shaevitz

## Abstract

Soil microbial communities regulate critical ecological processes, including nutrient cycling, carbon sequestration, and plant growth. However, due to the opacity and structural complexity of soil, how physical constraints imposed by pore geometry influence bacterial motility and chemotactic recruitment to plant roots remains poorly understood. We use a transparent soil mimic composed of cryolite grains that replicate the structural characteristics of natural soils while enabling direct visualization of bacterial dynamics. Using *Escherichia coli* as a model bacterium, we combine macroscopic spreading assays with microscopic tracking of cellular trajectories to characterize how soil texture affects motility across pore scales. We find that bacterial motility shifts from run-and-tumble behavior in large, open pores to frequent trapping in smaller, more confined spaces. This transition is governed by the pore size distribution and leads to reduced effective diffusivity and slower population-scale spreading. Moreover, pore-scale confinement hinders the chemotactic recruitment of bacteria to *Arabidopsis thaliana* roots: recruitment is robust in sandy and loamy soils but negligible in highly confining textures. Our results establish soil texture as a critical factor regulating microbial dynamics and ecological interactions in the rhizosphere. This mechanistic understanding complements genomic surveys by identifying physical confinement as an ecological filter that shapes root-associated microbiomes. These findings highlight the essential and previously underappreciated role of soil texture, suggesting new strategies for managing microbial communities to promote plant health and sustainable agriculture.

## Introduction

Soil harbors vast and diverse microbial communities that drive essential geological and biological processes, including nutrient cycling [1–3], carbon sequestration [4–6], and plant growth [7–9]. These microbial functions underpin ecosystem health, agricultural productivity, and the sustainability of terrestrial environments [10–13]. Central to many of these functions is microbial colonization of the rhizosphere, the soil region adjacent to plant roots, where bacteria influence plant physiology through nitrogen fixation [14], disease suppression [15], and other beneficial activities [16]. Importantly, the success of these interactions often depends on the ability of motile bacteria to locate and colonize root surfaces by chemotaxing toward exudates released into the surrounding soil [17, 18]. Indeed, rhizosphere environments tend to select for motile, flagellated bacteria, suggesting that movement and navigation play a central role in structuring root-associated microbiomes [19]. While recent genomic and metagenomic studies have expanded our understanding of microbial community composition and functional potential in heterogeneous soil environments [20–23], they provide limited insight into the physical processes that shape microbial distribution and function *in situ*. This gap is particularly relevant for agriculture, where understanding how bacteria locate and colonize plant roots could help guide the design of microbial-based alternatives to synthetic fertilizers and pesticides [24–27].

Our understanding of how the physical environment of soil impacts bacterial motility and rhizosphere colonization remains limited, due largely to the opacity and structural complexity of natural soils [6, 28, 29]. Most existing knowledge is based on genomic census data rather than direct mechanistic observation [2, 30–32]. A critical open question remains: how do microscale physical constraints imposed by soil pores modulate bacterial motility, spreading, and ultimately their ability to chemotax toward root exudates? Previous studies using transparent hydrogel packings have shown that bacterial motility can transition from run-and-tumble to intermittent hopping-and-trapping in confined geometries [33–37]. Run-and-tumble is the canonical paradigm in bulk liquids, consisting of straight “runs” punctuated by brief reorienting “tumbles” [38, 39]. In hopping-and-trapping, cells become transiently immobilized at constrictions and relocate via short “hops” between pores [33]. Microfluidic studies have also provided insight into bacterial motility [40] and navigation in chemical gradients, highlighting unique chemotactic responses in two-dimensional structured environments [41]. Whether two-dimensional approximations and the idealized geometries of soft bead packings adequately capture the structural complexity inherent to natural soils, however, remains unclear. Transparent soil analogues have enabled direct visualization of plant-microbe interactions in more soil-like conditions [42–46]. Most implementations emphasize bulk colonization or root-scale imaging rather than quantitative, pore-scale measurements of single-cell swimming and how those constraints propagate to population spreading and chemotactic recruitemnt at roots. Consequently, the effects of realistic soil textures on single-cell bacterial motility, multi-cellular migration, and recruitment to plant roots remain largely unexplored.

In this study, we use cryolite-based transparent soil microcosms that span textures from silt to sand, mimicking the pore size distributions of natural soils. These systems allow us to directly visualize and quantify how pore-scale confinement affects bacterial motility, spreading, and chemotaxis toward plant roots, using *Escherichia coli* and *Arabidopsis thaliana* as a model bacterium and plant, respectively. Our results reveal that bacterial motility transitions gradually rather than abruptly from run-and-tumble motion in open pores to frequent trapping in confining ones. This transition correlates with the abundance of pores smaller than the bacterial run length, reducing effective diffusivity and disrupting chemotactic recruitment. These results reveal how soil texture regulates microbial access to roots by modulating motility and chemotaxis, establishing a physical framework for engineering plant-associated microbiomes through soil structural manipulation.

## Results

### Transparent cryolite soils replicate realistic textures and enable pore-scale imaging

The opacity of natural soil presents a fundamental challenge to understanding soil microbial ecology, as it prevents direct visualization of microbial processes. To overcome this limitation, we use a transparent soil analogue composed of fragments of the mineral cryolite (Na_3_AlF_6_, sodium hexafluoroaluminate), which has a refractive index closely matching that of water (Fig. 1a) [47]. Although previous studies have used transparent soil analogues made from hydrogel or polymeric grains, we take advantage of cryolite to tune grain sizes and their distribution, creating more realistic granular textures while preserving optical clarity.

**Figure 1:**
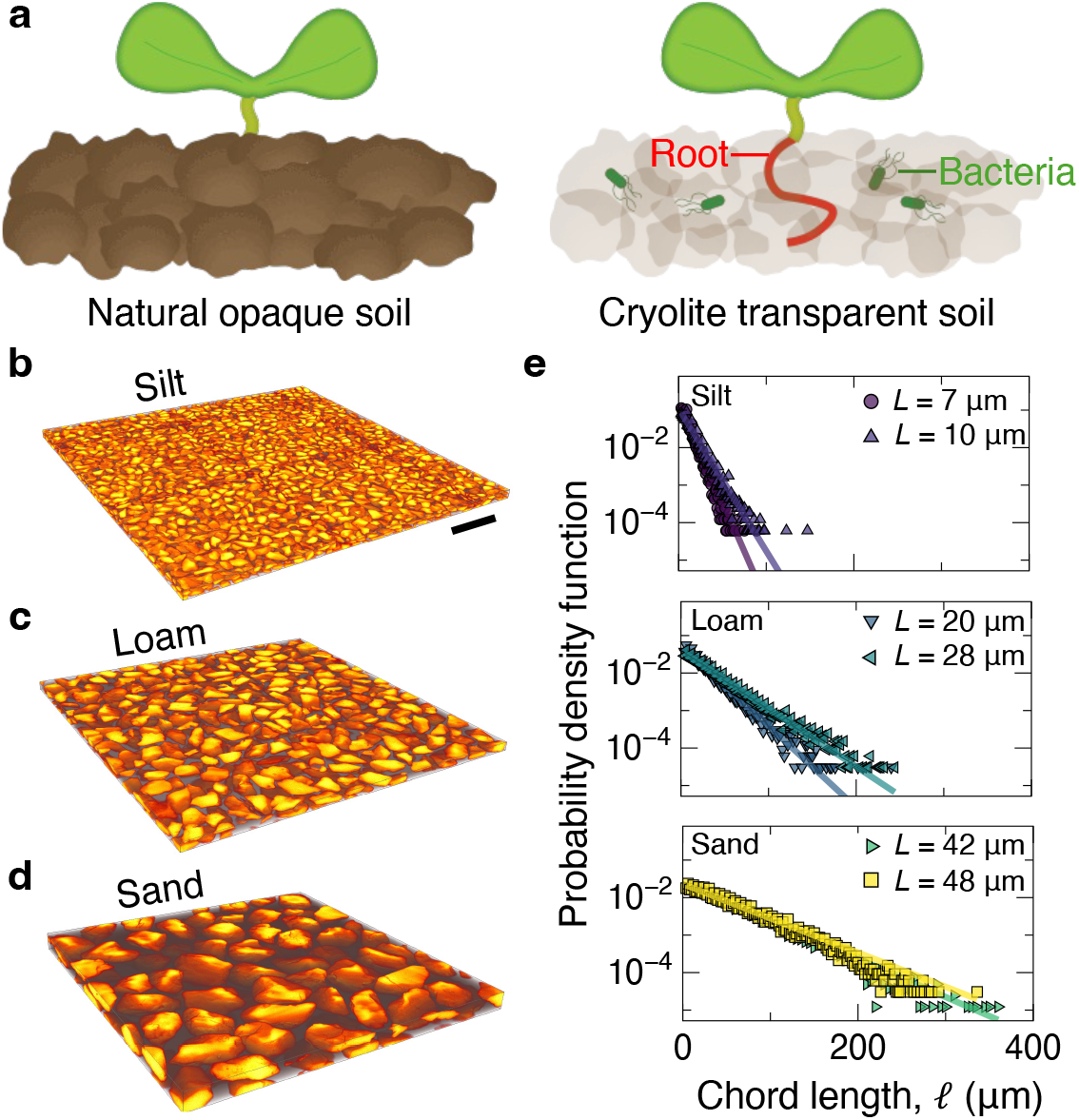
Cryolite-based transparent soil microcosms enable direct observation of bacterial dynamics and root interactions under controlled soil textures. (a) Schematic comparison between conventional opaque soil and transparent cryolite soil. (b–d) Confocal 3D renderings of cryolite packings for (b) silt, (c) loam, and (d) sand textures. Image intensity is inverted so that cryolite grains appear bright and pore spaces appear dark. Scale bar: 200 *µ*m. (e) Chord length distributions exhibiting exponential decay. Solid lines represent fits to Equation 1. *L* is the characteristic chord length.

Cryolite powder typically comes as a polydisperse mixture. To create transparent soil microcosms with well-defined textures, we separate the powder into six distinct particle size fractions, with two fractions each corresponding to silt, loam, and sand textures. Grain separation is achieved using a sonication-assisted wet sieving protocol that minimizes triboelectric aggregation of dry particles. Briefly, the polydisperse powder is placed on ASTM standard sieves immersed in an ultrasonic bath, while air is bubbled from beneath the mesh to continuously disrupt particle aggregates and facilitate efficient size separation. The sieve mesh sizes range from 32 *µ*m to 500 *µ*m, spanning the grain size distributions generally associated with silty, loamy, and sandy soils [48]. More details on transparent soil preparation are available in the Supplementary Information file (Note S1, Fig. S1, Table S1).

The optical clarity of saturated cryolite enables characterization of the resulting packings through direct imaging using confocal microscopy. We pack cryolite grains into wells of glass-bottom 24-well plates and saturate them with Berg’s Motility Buffer (BMB) containing 10 mM Rhodamine B dye to visualize the fluid-filled pore spaces, and 0.1 *µ*g mL^−1^ bovine serum albumin (BSA) to minimize hydrophobic interactions (Fig. 1b–d, Note S2). We reconstruct confocal image stacks to generate three-dimensional (3D) renderings of cryolite packings engineered to mimic realistic soil textures: silt (Fig. 1b), loam (Fig. 1c), and sand (Fig. 1d). Visual inspection of these 3D reconstructions shows that changes in particle size distribution strongly influence pore geometry, producing finer pore spaces in silty soils, larger interconnected pores in sandy textures, and a mixture of small and large pores in loamy environments.

To characterize these pore structures, we measure the chord length distribution of the pore spaces directly from confocal images. For each soil texture, we binarize pore space images and analyze them using a custom MATLAB script, sampling 1,000 random linear chords *𝓁* per image across multiple independent microscopy fields. The chord length distributions obtained for all soil textures exhibit exponential decay (Fig. 1e), consistent with theoretical expectations and empirical observations for random granular packings [49, 50]. This behavior arises from the statistical geometry of random packings, where the arrangement of irregularly sized grains naturally leads to a higher probability of short pore paths and an exponentially decreasing likelihood of encountering long, continuous pores.

We quantify the pore space distribution using a single characteristic length scale *L*, extracted by fitting the chord length probability density functions to an exponential decay model,

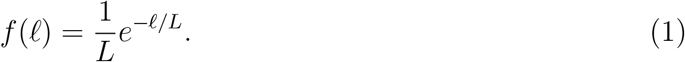

This characteristic length *L* serves as a meaningful metric for describing the dominant scale of pore confinement within each soil texture. More details on the characterization of soils can be found in Note S3.

### Soil texture modulates population-scale bacterial spreading dynamics

Having established the use of transparent cryolite soils with tunable pore size distributions, we next quantify how variations in soil texture modulate the ability of motile bacteria to spread within the porous medium. To do this, we directly observe the radial spreading dynamics of populations of fluorescently labeled, motile bacteria in cryolite soils saturated with BMB which contains no chemoattractants or nutrients; motion is therefore unbiased. Specifically, we use *Escherichia coli* constitutively expressing green fluorescent protein (GFP) as a model swimming bacterium and track its population-scale spreading behavior using time-lapse fluorescence confocal microscopy. We verified that cells remain motile in BMB over the duration of our experiments by measuring time-resolved velocity distributions up to 120 min (Fig. S2), corroborating previous evidence that *E. coli* swim for several hours in BMB by consuming endogenous energy reserves [51, 52]. We initiate the spreading experiments by injecting a small bacterial inoculum (2 *µ*L, ~ 10^9^ cells mL^−1^) into the center of wells containing packed cryolite grains saturated with media containing red Rhodamine B dye (Fig. 2a). Upon inoculation, the localized bacterial droplet forms a well-defined fluorescent bolus occupying a quasi-spherical volume of soil within the pore space (Fig. 2b). We then monitor bacterial spreading via time-lapse imaging at low magnification (2 × or 4 × objectives), enabling quantification of radial expansion dynamics over approximately 60 minutes post-injection (Fig. 2c).

**Figure 2:**
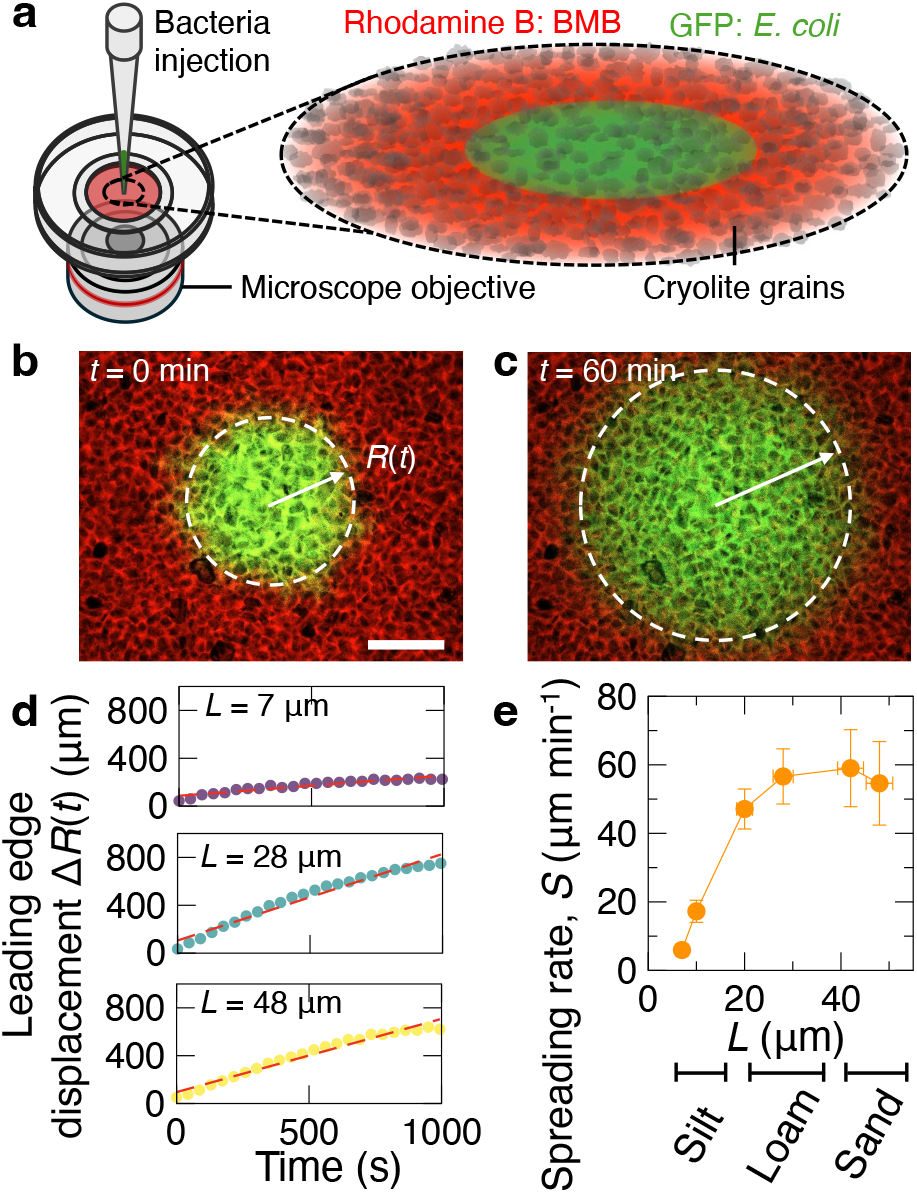
Effect of soil texture on the spreading dynamics of swimming bacterial populations. (a) Schematic illustration (left) of the experimental setup, in which a glass-bottom Petri dish containing a cryolite packing is imaged using an inverted fluorescence microscope. Bacteria are introduced locally into the porous medium via micropipette injection. Inset (right) depicts a zoomed schematic of the cryolite packing saturated with Berg’s Motility Buffer (BMB; red fluorescence due to Rhodamine B) and a localized inoculum of GFP-expressing *Escherichia coli* (green). (b–c) Representative confocal microscopy images showing bacterial spreading within the porous medium at (b) initial injection (0 min) and (c) after 60 minutes of spreading. The dashed white circle indicates the edge of the bacterial population, *R*(*t*), highlighting radial expansion over time. Scale bar: 1 mm. (d) Quantification of bacterial population spreading over time, showing displacement of the leading edge, Δ*R*(*t*), for soil textures representing silt (*L* = 7 *µ*m; top), loam (*L* = 28 *µ*m; middle), and sand (*L* = 48 *µ*m; bottom). The red dashed line indicates the linear fit used to determine the spreading rate *S*. (e) Spreading rate *S* of bacterial populations as a function of characteristic pore length *L*. Bacteria spread readily in sandy and loamy textures but are significantly hindered in silty textures. Horizontal error bars represent the 95% confidence interval for *L*, obtained from exponential fits to the chord length distributions in Fig. 1. Vertical error bars represent the standard deviation across *n* = 3 biological replicates.

We observe that bacterial spreading within the porous media is consistently subdiffusive, as evidenced by analyzing the radial spreading profiles (see Supplementary Information for details, Note S4, Figures S2-3). Due to the subdiffusive nature of the spreading, extracting a diffusivity parameter is problematic and potentially misleading. Therefore, we opt for a physically meaningful measure of bacterial transport by tracking the leading edge of the spreading front, *R*(*t*). Specifically, we approximate the bacterial expansion as linear over the initial 1000 s time interval, allowing us to quantify the spreading dynamics by defining a spreading rate, *S*, as the slope of the leading edge displacement, Δ*R*(*t*), during this initial linear regime. To do this, we implement an image analysis protocol that extracts *R*(*t*) from fluorescence intensity profiles obtained from image sequences. Radial intensity profiles are computed by azimuthal averaging of pixel intensities over concentric annuli centered on the inoculum. *R*(*t*) is identified as the radial location where fluorescence intensity exceeds a threshold defined as 5% above background, based on regions well outside the initial bacterial bolus.

Representative data for the leading edge displacement, Δ*R*(*t*) = *R*(*t*) − *R*(0), across the three soil textures highlight significant differences in bacterial spreading dynamics as a function of pore geometry (Fig. 2d). Bacteria spread rapidly in porous media representative of sandy and loamy textures, characterized by large pores (20 *µ*m *< L <*50 *µ*m), with spreading rates of approximately 50–60 *µ*m (Fig. 2e). By contrast, spreading is hindered in soils representing silty textures with smaller, more confining pore structures (*L <* 10 *µ*m), where rates decrease by roughly an order of magnitude to 5–10 *µ*m min^−1^. This sharp, nonlinear decline in spreading rate at small pore sizes suggests a strong dependence of bacterial motility and transport on pore geometry.

### Pore-scale analysis reveals a transition from run-and-tumble motility to trapping dictated by pore geometry

To explain differences in bacterial spreading across soil textures, we examine how pore-scale confinement affects motility at the single-cell level. We hypothesize that soil pore geometry alters bacterial movement by inducing a transition from run-and-tumble dynamics in open pores to constrained behavior in tighter spaces, similar to the hopping-and-trapping seen in hydrogel systems [33–35] (Fig. 3a). Because natural soils and our cryolite packings have broader pore size distributions and more irregular geometries, the resulting dynamics may be more complex than those observed in hydrogels.

**Figure 3:**
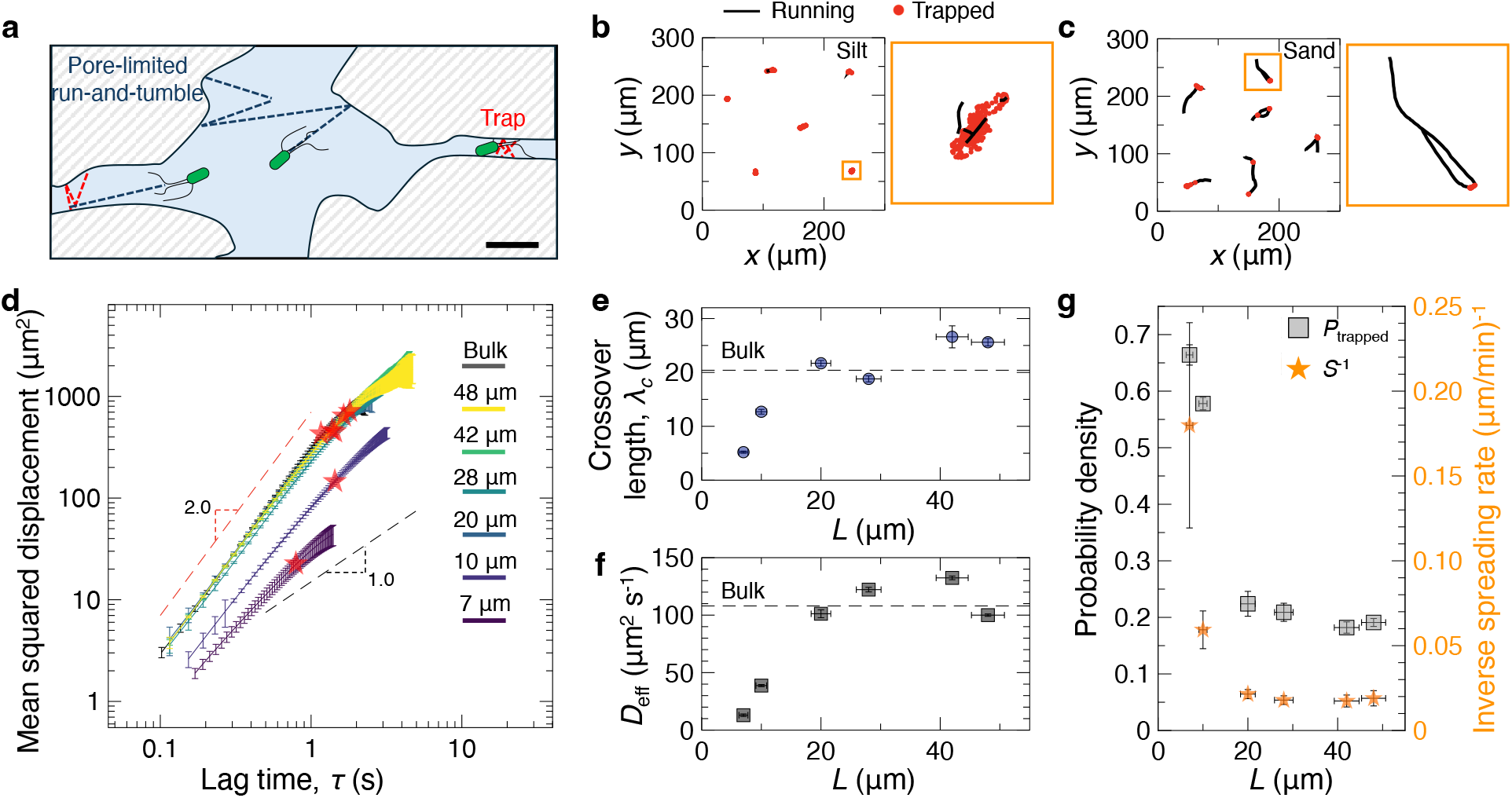
Pore-scale structure dictates the transition from bacterial running to trapping. (a) Schematic illustrating pore-limited bacterial dynamics in soil, showing transitions from run-and-tumble motility in open pores (black dashed lines) to extended trapping events in highly confining pores (red lines). (b–c) Representative trajectories of bacteria in cryolite packings mimicking (b) silt and (c) sand textures. Trajectories are 5 ± 0.5 s in duration; orange-outlined panels show magnified views of the boxed regions, highlighting a single representative trajectory. Black lines denote running states, and red circles indicate trapped states, as classified in the Supplementary Information. (d) Ensemble-averaged mean squared displacement (MSD) curves for bacterial populations in environments with different characteristic pore lengths *L* (legend). MSDs show ballistic (∝ *τ* ^2^) behavior at short lag times and transition to diffusive (∝ *τ*) behavior at longer lag times; red stars mark the crossover points. Dashed lines indicate slopes corresponding to ballistic (2.0) and diffusive (1.0) regimes. (e) Crossover lengths extracted from the MSDs as a function of *L*, illustrating that the length scale of persistent motion increases with pore size and plateaus at the bulk value for *L >* 20 *µ*m (dashed line: bulk fluid). (f) Effective diffusivity (*D*_eff_) as a function of *L*, increasing with pore size and approaching the bulk diffusivity ( ~ 100 *µ*m^2^ s^−1^, dashed line) for large *L*. (g) Correlation analysis: the probability of a bacterium being trapped (*P*_trapped_, squares) decreases monotonically with increasing pore size, and inversely tracks the population-scale spreading rate (orange stars). Error bars in (d) represent the standard error of the mean across all tracks available for each lag time. Horizontal error bars in panels (e), (f), and (g) represent the 95% confidence interval from exponential fits to the pore chord length distributions (see Fig. 1). Vertical error bars in (e) and (f) are the 95% confidence intervals from linear fits to the MSD analysis (see Methods). For *P*_trapped_ in (g), error bars show the standard error of the mean from all frames pooled across *>*15,000 frames per condition (minimum 300 tracks). Vertical error bars for the inverse spreading rate are standard deviations from three biological replicates.

To test this, we perform direct pore-scale imaging of bacterial trajectories within the cryolite microcosms and analyze motility on a frame-by-frame basis. Each frame in a trajectory is classified as either running or trapped using a custom algorithm. We first compute a displacement ratio *δ*(*t*) for each frame, defined as the net displacement over a temporal window, *w* = 20 frames (or 0.8 s), divided by the total path length traveled during that interval. This ratio serves as an inverse measure of local tortuosity: high values indicate directed motion, while low values reflect localized movement with minimal net displacement. Frames with *δ*(*t*) *<* 0.6 are classified as trapped.

For frames not classified as trapped by the displacement ratio, we apply additional criteria based on instantaneous velocity and angular reorientation to classify runs. A frame is classified as running only if the velocity exceeds 10 *µ*m s^−1^ and the directional change is less than *π/*6. Frames failing these thresholds are reclassified as trapped. Thresholds for *δ*(*t*), velocity, and angular change were validated against manually labeled trajectories and optimized for precision and recall to ensure consistent and reproducible classification (see Supplementary Information for details, Note S5 and Fig. S5).

Applying this classification across a large set of trajectories reveals that bacteria in soil frequently exhibit mixed motility within a single trajectory. Individual cells often alternate between run-and-tumble motion and transient trapping as they navigate the heterogeneous pore network. This behavior contrasts with that observed in hydrogel packings that contain a single pore lengthscale, where motion is characterized by abrupt transitions between fully trapped states via discrete hops through narrow constrictions [33–35]. The coexistence of motility modes in soil reflects the broad distribution of pore sizes inherent to realistic granular media.

Analysis of the classified trajectories shows a wide distribution of bacterial run lengths (*L*_run_) across soil textures. Rather than exhibiting a clear characteristic length scale, run lengths vary continuously from short excursions limited by pore throats of about 2-5 *µ*m (Fig. 3b) to longer trajectories of 40-50 *µ*m in loamy and sandy soils (Fig. 3c, Fig. S6-7). While this broad distribution prevents straightforward quantitative interpretation, it directly reflects the structural complexity and heterogeneity of realistic soil pore networks. In addition, these observations further highlight that bacteria in soil experience varied patterns of motility, reinforcing the need to fully characterize the impact of pore-scale confinement on bacterial swimming.

### Mean squared displacement reveals how pore-scale confinement shapes population spreading

Given the absence of clear characteristic scales from run length distributions alone, we quantify the effects of confinement more rigorously by applying a mean squared displacement (MSD) analysis to individual bacterial trajectories. MSD provides a well-established framework for characterizing microscopic dynamics, including run-and-tumble motion, by distinguishing between regimes of movement [53]. At short lag times *τ*, ballistic motion is indicated by quadratic scaling (MSD ∝ *τ* ^2^), while at longer times, diffusive behavior emerges with linear scaling (MSD ∝ *τ*). These distinct scaling regimes reflect the underlying physical processes. Ballistic behavior at short *τ* corresponds to persistent swimming during runs, whereas diffusive dynamics at longer *τ* arise from directional reorientations due to tumbling, collisions with soil grains, and other confinement effects.

We compute the ensemble-averaged MSD, ⟨MSD(*τ*)⟩, by first tracking individual bacteria over thousands of frames and calculating each trajectory’s MSD, defined as

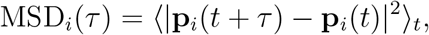

where **p**_*i*_(*t*) is the position of bacterium *i* at frame *t*, and the average is taken over all timepoints *t*. We then average across all trajectories to obtain a statistically robust ensemble MSD. More details on the analysis are available in Note S6.

Across all soil textures, the ensemble MSD exhibits a consistent transition from ballistic behavior at short lag times to an approximately linear, diffusive behavior at longer lag times (Fig. 3d). We identify this crossover using segmented regression on log-log plots of the MSD curves. Two linear segments, representing the ballistic and diffusive regimes, are fit simultaneously. Their intersection, (*λ*_*c*_, *τ*_*c*_), defines the crossover point (red stars in Fig. 3d). Physically, this point marks the spatiotemporal scale at which bacteria shift from persistent swimming to randomized motion due to tumbling, pore collisions, and trapping. Notably, this linear regime in the MSD probes motility at lag times of a few seconds that lie between the ballistic regime and the minute-to-hour scales at which population-level assays reveal subdiffusive spreading.

We find that the measured crossover length *λ*_*c*_ increases systematically with the characteristic pore length *L* (Fig. 3e). In soils with small pores (*L <* 20*µ*m), *λ*_*c*_ increases approximately linearly with *L*, indicating that pore geometry directly constrains the extent of ballistic motion. In this regime, bacteria typically reach a pore boundary before they tumble, so the transition from ballistic to diffusive behavior is set by physical confinement. In contrast, for soils with larger pores (*L >* 20 *µ*m), *λ*_*c*_ reaches a plateau around 25–30 *µ*m, even though pore sizes continue to increase (*L* = 42–48 *µ*m in sandy soils). This plateau matches the crossover length measured in bulk media, where motion is unconfined and the ballistic-to-diffusive transition arises from intrinsic motility processes such as run-tumble dynamics and rotational diffusion. Thus, the scaling of *λ*_*c*_ with *L* reveals a shift from boundary-limited motion in small pores to motility-limited dynamics in large pores.

At the lag times where diffusive behavior emerges, the ensemble MSD provides a direct estimate of an effective diffusivity, defined for two-dimensional measurements as *D*_eff_ ≡ ⟨MSD(*τ*)⟩*/*4*τ*. We extract *D*_eff_ from linear fits to the MSD curves in the diffusive regime for each soil texture (Fig. 3f). Consistent with the variation of *λ*_*c*_, *D*_eff_ varies strongly with pore geometry as well. In silty soils with small characteristic pores, *D*_eff_ is sharply reduced, with values as low as approximately 10 *µ*m^2^s^−1^. In contrast, for loamy and sandy soils with larger pores, *D*_eff_ approaches approximately 100 *µ*m^2^ s^−1^, matching values measured in bulk fluid. We emphasize that *D*_eff_ is an operational, scale-specific mobility extracted from the MSD’s linear regime and is distinct from the population-level subdiffusion characterized by the spreading rate *S* in Fig. 2. The reduction in diffusivity in fine-textured soils directly quantifies how pore-scale confinement limits bacterial exploration and diminishes population-scale spreading.

Ultimately, the connection between pore-scale confinement, bacterial run-tumble-trap dynamics, and population-level spreading depends on the trapping rate of cells. We therefore quantify the fraction of time cells spend in trapped states, *P*_trapped_, identified from trajectory analysis across the various soils. This measure varies with soil texture showing that tighter pore geometries impose more frequent and longer-lived physical constraints on swimming cells.

To connect these microscopic constraints to macroscopic behavior, we examine how *P*_trapped_ relates to the bacterial spreading rate. The increase in trapping time with decreasing *L* mirrors the inverse of the bulk spreading rate: soils in which cells spend a larger fraction of time trapped correspondingly slow the advance of the population from the initial inoculum (Fig. 3g). This correspondence indicates that the soil texture governs how fast populations of bacteria migrate by way of modulating the rate of trapping. Under progressive confinement, bacteria do not switch between two distinct motility modes, but instead exhibit continuously shorter effective runs and elevated trapping, which together set the macroscopic transport speed.

### Soil texture influences bacterial chemotaxis toward plant roots

The results presented so far demonstrate how soil pore geometry influences the run-and-tumble motility of bacteria. However, in natural ecosystems, bacterial movement is often not random but directed by chemical gradients, i.e. via chemotaxis, toward sources of nutrients or signaling molecules [54, 55]. In soil, plant roots secrete a diverse array of chemical compounds known as root exudates, which can serve as chemoattractants drawing microbial populations toward the root surface [56]. However, the influence of physical confinement within realistic soil textures on these chemically driven root–microbe interactions remains poorly understood [57, 58]. Given our findings on the regulation of bacterial motility by soil texture, we hypothesize that such physical constraints also modulate chemotactic recruitment toward plant roots.

To test this hypothesis, we perform controlled chemotactic assays by introducing living plant roots into the microcosms in the presence of homogeneously distributed bacteria. Specifically, we use roots of *Arabidopsis thaliana*, a model plant extensively studied in microbiome research and recently introduced into hydrogel-based porous media for root phenotyping [59]. Sterilized *A. thaliana* seeds are initially germinated and grown in ultrapure water to minimize contamination and residual nutrients. We then fluorescently label post-germination seedlings by incubating them with Rhodamine B dye. The seedlings are carefully transplanted into cryolite soil microcosms, which are then saturated with BMB containing *E. coli*. Although *E. coli* is not traditionally considered a rhizosphere bacterium, its chemotaxis system is tuned to small metabolites that are abundant in plant root exudates [60]. These include amino acids such as glycine, alanine, and isoleucine, or simple sugars such as galactose and glucose, and dipeptides [61, 62], which are known *E. coli* chemoeffectors [63, 64]. Consistent with this mechanisms, *E. coli* was previously shown to exhibit chemotaxis toward plant tissue extracts, and mutants defective in chemotaxis are severely impaired in their ability to colonize plant roots and leaves [65]. Therefore, we observe the chemotactic interaction between flagellated bacteria and plant roots using *E. coli* at an initial concentration of approximately 1 × 10^7^ cells mL^−1^. We directly visualize and quantify bacterial movement and accumulation near the root surface over approximately 60 minutes (Fig. 4a–c).

**Figure 4:**
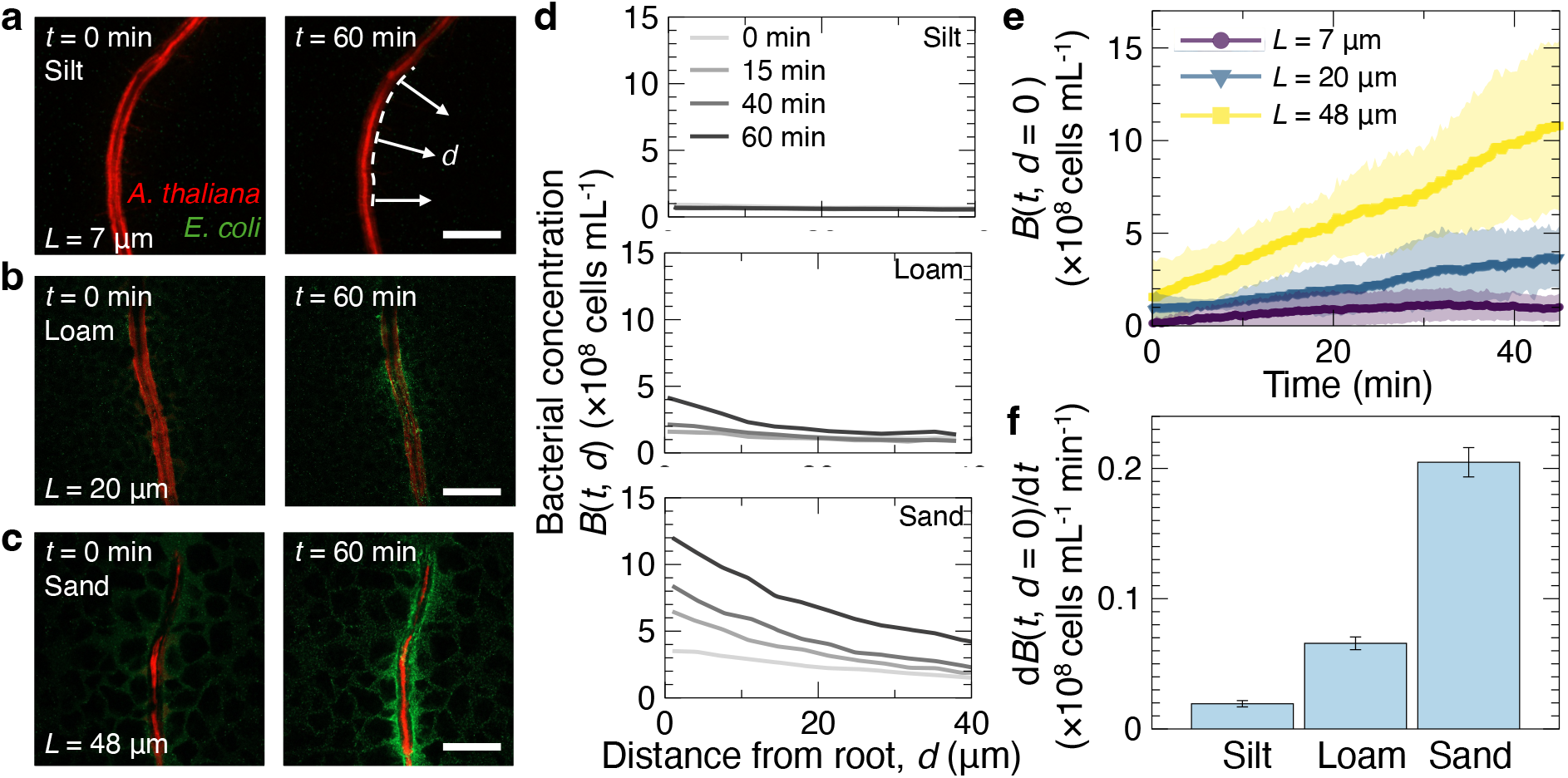
Impact of soil texture on root–bacteria interactions mediated by bacterial motility. (a–c) Representative confocal microscopy images showing *Arabidopsis thaliana* roots (red, labeled with Rhodamine B) interacting with GFP-expressing *E. coli* (green), initially suspended at approximately 1 × 10^7^ cells mL^−1^. Images represent (a) silt, (b) loam, and (c) sand textures. Left panels: initial conditions at inoculation (*t* = 0 min). Right panels: bacterial accumulation after 60 minutes. Bacterial concentration is quantified as a function of distance *d* from the root surface. Scale bar: 100 *µ*m. (d) Radial bacterial concentration profiles *B*(*t, d*), averaged over 100 sampling lines perpendicular to the root. Accumulation is minimal in silt (top), moderate in loam (middle), and strong in sand (bottom). (e) Temporal evolution of bacterial concentration at the root interface (*d* = 0), averaged over three biological replicates for textures with *L* = 7 *µ*m (silt), *L* = 20 *µ*m (loam), and *L* = 48 *µ*m (sand). Shaded regions indicate standard deviation across three replicates. (f) Accumulation rate at the root interface (*dB*(*t, d* = 0)*/dt*) for each soil texture. Bars represent the mean of three biological replicates with error bars indicating the standard error. One-way ANOVA showed a significant effect of soil texture (*p* = 0.031).

In a typical experiment, bacteria are initially distributed uniformly throughout the fluid-filled pore spaces, with no preferential localization near the roots. Over time, we observe texture-dependent bacterial accumulation at the root surface. In sandy soils, bacteria accumulate rapidly, forming distinct layers adjacent to the root within tens of minutes. A similar, though slower, accumulation is observed in loamy textures. In silty soils, bacterial accumulation is minimal or absent, despite the presence of root-derived chemical attractants. To verify that this absence was not simply due to complete suppression of exudate transport, we measured the spreading rate of Rhodamine B—which we use as a proxy for the small molecule metabolites in root exudates— in each soil (see Fig. S8 in Supplementary Information file). Although Rhodamine B transport slows by approximately two-fold in silt compared to sand, declining in a roughly linear fashion with mean pore size, gradient formation still occurs. Overall, these observations suggest that chemotactic signaling between roots and bacteria is strongly modulated by soil texture not purely due to compression of the exudate gradients, but also due to the role of confinement on bacterial motion.

We quantitatively assess bacterial recruitment by measuring the distribution of bacteria as a function of distance *d* from the root surface, i.e., the rhizoplane. Briefly, we segment the root images to extract root shape and skeletonize the resultant mask to compute a smooth centerline. At each point along the centerline, we construct normal vectors to define a systematic sampling grid (Fig. S9). Using this grid, we extract fluorescence intensity profiles of bacteria along lines emanating from the root surface (with *d* = 0 at the rhizoplane), which reflect local bacterial concentration. We then average these profiles longitudinally, and calibrate the fluorescence intensity to cell density using independent colony-forming unit (CFU) measurements (Fig. S10, Fig. 4d). From these calibrated profiles, we evaluate bacterial concentration at the root interface (*d* = 0) to quantify the temporal evolution of root-associated accumulation (Fig. 4e).

Consistent with our qualitative observations, root-adjacent bacterial concentration varies markedly with soil texture. In sandy soils, accumulation increases linearly over time, reaching levels significantly higher than in loam or silt (Fig. 4e). Loamy soils exhibit intermediate accumulation, while silty soils show negligible recruitment, highlighting the strong constraint imposed by fine-scale physical confinement.

Through a linear fit to the bacterial concentration at the root surface over time, we extract the accumulation rates as a measure of bacterial recruitment efficiency (Fig. 4f). These rates reveal clear differences across soil textures: bacterial accumulation at roots occur slowly in silty roots, at a rate of (0.019 ± 0.002) × 10^8^ cells mL^−1^ min^−1^, increases substantially in loamy soils to (0.066 ± 0.005) × 10^8^ cells mL^−1^ min^−1^, and reaches a maximum in sandy soils at (0.20 ± 0.01) × 10^8^ cells mL^−1^ min^−1^. In hydroponic controls (roots without cryolite), colonization proceeds too rapidly to quantify under the same acquisition settings: bacteria attach and the fluorescence signal increases rapidly in the first minutes after inoculation, leaving nearly no linear window for rate estimation. While restricting quantitative comparisons to soil, we can note that even the least confining sandy texture slows recruitment substantially *t* relative to hydroponics. These results highlight a critical, previously unmeasured role of soil physical matrix in mediating chemically driven interactions between roots and microbes. Specifically, the measurement reveals a new mechanistic insight into root-bacteria interactions in which chemical signaling through root exudates alone is insufficient to guarantee bacterial recruitment under realistic, physically heterogeneous soil conditions. Instead, the physical constraints imposed by soil pore geometry play a decisive role in modulating and in extreme cases overriding the chemotactic response.

## Discussion

The opacity of natural soil has long impeded efforts to understand microbial dynamics and interactions by preventing direct observation of behavior *in situ*. While genomic and metagenomic approaches have greatly expanded our knowledge of microbial diversity and ecological function in soils, they primarily yield census-based snapshots rather than mechanistic insights into underlying processes [66]. By leveraging transparent cryolite soil microcosms, we addressed this gap with a platform that replicates the heterogeneity and complexity of natural soil textures while allowing direct visualization of bacterial motility at both pore and population scales. Our results show that bacterial transport is strongly regulated by soil texture: populations spread rapidly in sandy and loamy textures but are substantially hindered in silty soils. Through direct imaging of bacterial trajectories, we find that these macroscale differences arise from continuous changes in motility patterns, not discrete transitions. Rather than switching abruptly from run-and-tumble to hopping-and-trapping behavior, bacteria in granular cryolite frequently alternate between running, tumbling, and transient trapping, even within single trajectories. This behavior emerges naturally from the broad pore size distributions characteristic of realistic soil environments.

To quantitatively connect microscale motility to macroscale spreading, we applied mean squared displacement analysis. Across soil textures, we observed a clear ballistic-to-diffusive crossover, with the crossover length increasing with the characteristic pore size *L*. In soils with larger pores, motility closely resembles the behavior seen in liquid, with crossover lengths approaching those measured under unconfined conditions. In contrast, silty soils curtail ballistic runs and reduce bacterial diffusivity by roughly an order of magnitude. Notably, both the probability of trapping and the fraction of pores smaller than the average bacterial run length are strongly inversely correlated with spreading rate. These findings show that geometric confinement at the pore scale exerts upward control over population-scale dynamics, modulating bacterial transport through its effects on motility.

Beyond intrinsic motility, these effects have direct implications for ecological interactions driven by chemotaxis. Chemotaxis underpins a broad range of microbial behaviors relevant to nutrient acquisition, symbiosis, and host colonization. Our experiments incorporating living *A. thaliana* roots provide direct evidence that soil structure can override chemotactic signaling mediated by root exudates. Roots embedded in coarse-textured soils robustly recruit bacteria, while those in silty soils show negligible bacterial accumulation despite comparable chemical cues. These results reveal that pore-scale confinement, rather than being a passive background, can actively modulate or even suppress microbial recruitment by limiting the physical feasibility of chemotactic navigation. The strong texture-dependent modulation of bacterial-root interactions demonstrates that physical constraints can dominate over chemical signaling pathways traditionally assumed sufficient for recruitment and assembly.

These findings could have far-reaching ecological implications. Because motility and chemotaxis are central to microbial interactions in soils, the heterogeneity of natural textures likely acts as an ecological filter, selectively permitting or restricting colonization based on motility characteristics. While our experiments used model microswimmers, the same principles are likely to apply to other flagellated bacteria, including plant symbionts such as *Rhizobia* and pathogens such as *Pseudomonas*. The large differences in recruitment rates across textures suggest that soil physical structure could influence plant health and disease dynamics by shaping root-associated microbiomes. These insights also hold relevance for sustainable agriculture, where microbial-based approaches increasingly seek to replace chemical inputs with beneficial microbes. The success of such strategies depends on effective colonization of roots, which we find to be strongly constrained by soil texture. Understanding how pore-scale structure shapes these interactions opens new opportunities for managing soil physical properties to improve microbial recruitment and optimize plant–microbe interactions in realistic agricultural settings.

## Materials and Methods

### Materials

Cryolite powder was obtained from Andover Corporation. Luria-Bertani (LB) broth and agar for bacterial culture, along with potassium phosphate dibasic (K_2_HPO_4_), potassium phosphate monobasic (KH_2_PO_4_), sodium chloride (NaCl), ethylenediaminetetraacetic acid (EDTA), Rhodamine B dye, and bovine serum albumin were all purchased from Sigma-Aldrich. Glass-bottom Petri dishes and well plates used for microscopy experiments were purchased from Cellvis. The list of ASTM standard separation sieves used (Gilson Inc.) is available in the Supplementary Information.

### Bacterial culture

We use *E. coli* strain W3110, which constitutively express GFP throughout the cytoplasm. To prepare a liquid bacterial culture, a single colony from an LB agar plate is inoculated into 2 mL of LB broth. Cultures are grown overnight at 30^°^C with shaking. The following day, the overnight culture is diluted 1:100 into fresh LB medium and incubated at 30^°^C with shaking for an additional 3 hours before use in experiments.

### Plant growth

We use *A. thaliana* ecotype Columbia-0 (Col-0) seeds. Seeds are sterilized by immersing them in 70% ethanol in a vial and gently rotating for 5 minutes [67]. The ethanol is removed by pipetting, and this washing step is repeated with fresh ethanol a total of three times. The sterilized seeds are then dried aseptically in a biosafety cabinet and stored under sterile conditions. To initiate plant growth, sterilized seeds are individually placed into separate wells of a 96-well plate containing autoclaved DI water. Seeds undergo stratification at 4^°^C in darkness for two days. Following stratification, seeds are transferred to room temperature under constant light for germination. Germination typically takes three days, and seedlings are used within two days post-germination for experiments. Prior to experiments, seedlings are stained by incubating them in a 50 mM Rhodamine B solution for 10 minutes. Excess dye is removed by gentle pipetting, and seedlings are rinsed multiple times with fresh sterile DI water to eliminate residual dye.

### Imaging and tracking

All imaging was performed on a Nikon A1R inverted confocal microscope (Eclipse Ti2), using 488 nm and 561 nm wavelength lasers. Galvano scanning was used for single images, z-stacks, and slow timelapses (up to 5 fps), and resonant scanning was used for fast timelapses (up to 50 fps). Objectives used included 2 × (0.1 N.A.), 4 × (0.2 N.A.), 10 × (0.3 N.A.), and 20 × (0.8 N.A.), all dry. Additional imaging parameters and acquisition settings are provided in the Supplementary Information. Individual bacteria were tracked using the TrackMate plugin in ImageJ. Bacteria were first segmented as spots in each frame using a Laplacian-of-Gaussian filter with sub-pixel localization. The plugin then linked detected spots between consecutive frames by solving a linear assignment problem that minimizes the cost across all possible spot-to-spot connections.

## Acknowledgments

We thank the members of the Datta and Shaevitz groups, as well as Prof. Victoria Orphan for useful discussions. We also thank Dr. Meera Ramaswamy for assistance with CFU measurements; Ella Gunady and Prof. Jonathan Conway for providing *A. thaliana* seeds; and Dr. Michael Bunsick for help with plant protocols. AAH and JWS acknowledge support from the Center for the Physics of Biological Function (CPBF) and the National Science Foundation (NSF PHY-1734030). We acknowledge support to SSD from the National Science Foundation (NSF CBET-1941716) as well as the Camille Dreyfus Teacher-Scholar and Pew Biomedical Scholars Programs.

## Author contributions

A.A.H., S.S.D., and J.W.S. designed research. A.A.H. and G.C. performed research. S.S.D. and J.W.S. contributed new reagents/analytic tools. A.A.H., S.S.D., and J.W.S wrote the paper.

## Supplementary Information

### Note S1: Cryolite processing and soil sample preparation

We obtain pre-milled natural cryolite powder with a broad initial grain size distribution (Andover Inc.). The particle sizes range from sub-50 *µ*m to several hundred microns. We prepare transparent soils with tunable textures by fractionating this powder into narrower size ranges using the following wet sieving protocol. The fractionation setup is illustrated in Figure S1. We place cryolite, typically in 5 g aliquots, on ASTM standard brass sieves (Gilson Inc.; see Table S1 for specifications) and immerse the sieves in a beaker filled with ultrapure Milli-Q water and equipped with a tube connected to the laboratory pressurized air source. We place this beaker in an ultrasonication bath and introduce vertical bubbling from the tube positioned below the sieve. The combination of continuous air bubbling and ultrasonic agitation disperses agglomerates and enhances particle mobility across the sieve mesh. This facilitates the passage of undersize grains while avoiding potential triboelectric aggregation. We perform each sieving step for 10 minutes under these standardized conditions for all sieve sizes. After size separation, we collect each cryolite fraction in water and sterilize it through a multistep solvent exchange. First, we pellet the particles by centrifugation at 3000 rpm for 1 minute in a standard benchtop centrifuge. Then, we resuspend the pellet in 50 mL of absolute ethanol, vortex briefly, and gently rotate for 10 minutes. We repeat this step twice and follow it with one rinse in 70% ethanol and one rinse in autoclaved Milli-Q water. After the final rinse, we air dry the cryolite overnight in a biosafety cabinet before storing the dry powder. For imaging, we load ~ 500 mg of cryolite either into 35 mm glass-bottom dishes or individual wells of 24-well glass-bottom plates. We saturate the cryolite packings with Berg Motility Buffer (BMB; 6.2 mM K_2_HPO_4_, 3.8 mM KH_2_PO_4_, 67 mM NaCl, 0.1 mM EDTA) [**?**] supplemented with 10 mM rhodamine B (Sigma-Aldrich) and 0.1 *µ*g mL^−1^ bovine serum albumin (BSA; Sigma-Aldrich). Rhodamine B uniformly stains the aqueous phase for clear detection of the pore space, and BSA minimizes nonspecific adsorption to cryolite surfaces.

**Figure S1:**
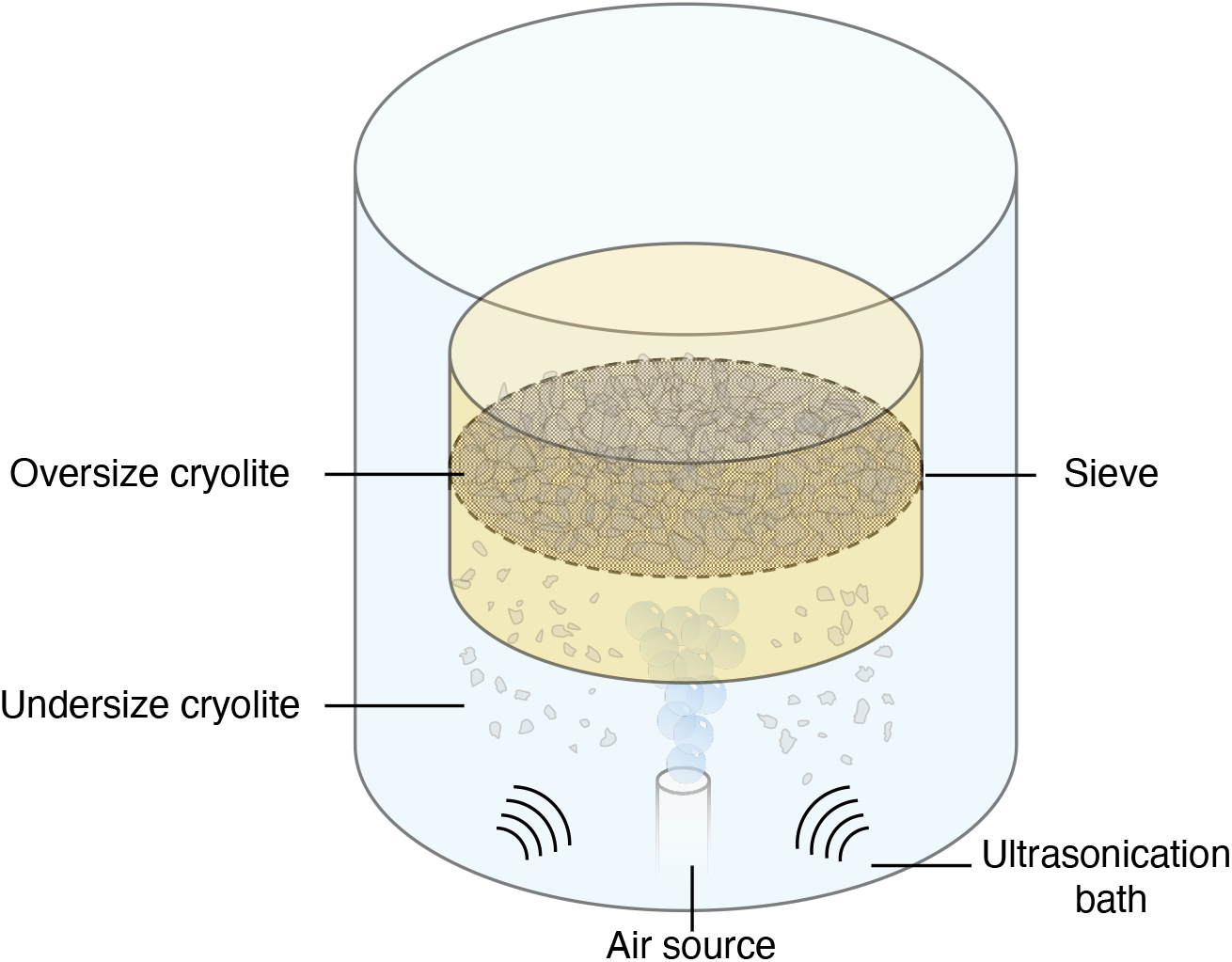
Schematic of the cryolite wet sieving setup. We place the cryolite powder on an ASTM standard sieve submerged in ultrapure water. An air source positioned below the sieve introduces vertical bubbling while an ultrasonication bath applies acoustic agitation. This disperses clump and enhances separation of undersize and oversize cryolite particles.

**Table S1:** ASTM standard brass sieves used for cryolite size fractionation. Each sieve is half-height and 3 inches in diameter.

| Sieve ASTM designation | Mesh size |
| --- | --- |
| 450 | 32 $\mu\text{m}$ |
| 270 | 53 $\mu\text{m}$ |
| 200 | 75 $\mu\text{m}$ |
| 140 | 106 $\mu\text{m}$ |
| 100 | 150 $\mu\text{m}$ |
| 35 | 500 $\mu\text{m}$ |

### Note S2: 3D renderings of cryolite packings

We generated three-dimensional renderings of the cryolite transparent soils using a Nikon AX confocal microscope equipped with a Ti2 stand and a galvano scanner. We used a PLAN APO *λ* 20× objective with a numerical aperture of 0.8 and excited the rhodamine B dye in the aqueous phase with a 561 nm laser and detected emission using a 595/50 nm bandpass filter. We acquired *z*-stacks with a step size of 0.975 *µ*m, covering at least 50 *µ*m vertical depths while stitching multiple *xy* fields of view to obtain images covering at least 1 × 1 mm areas. Note that we limited imaging depth to approximately 300–400 *µ*m to minimize optical aberrations due to compounding small mismatches in refractive index. We processed the image stacks in *ImageJ* to crop the field of view and invert the LUT to visualize the grain space as opposed to the pore space. Finally, we generated 3D renderings using the volume viewer toolset built into the Nikon ND software.

### Note S3: Soil characterization via chord length distribution

We characterized pore geometry in cryolite transparent soils by analyzing chord length distributions extracted from 2D fluorescence confocal slices. We selected slices taken approximately 100 *µ*m above the glass bottom of the dish to avoid packing artifacts caused by surface effects. We binarized each frame using Otsu’s thresholding method to separate the fluorescent aqueous phase representing pores (assigned a value of 1) and the cryolite grain phase (assigned a value of 0). In each binary frame, we superimposed 1000 randomly positioned lines and identified contiguous runs of constant pixel value along each line. These uninterrupted segments are the chords *𝓁*, defined as segments lying entirely within one phase and bounded by an interface with the opposite phase. We computed the length of each chord while excluding all segments shorter than 2 pixels ( ~ 1.7 *µ*m) to avoid artificial inflation of short-length counts. We repeated this procedure across 10 independent frames per soil condition, aggregating chord data for the pore phase. We fitted the resulting chord length distribution *p*(*𝓁*) with a single-parameter exponential decay model, 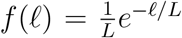. Here, the exponential decay parameter *L* is the characteristic length that we interpret as the representative geometric scale of the pore phase. We used nonlinear least squares to fit this model to the pore chord distributions and report the decay length as a quantitative descriptor of soil pore geometry. This method is supported by prior work on statistically isotropic and uncorrelated random media, where chord length distributions asymptotically decay exponentially in the absence of long-range order. In our data, this model captures the observed distributions well across all soil conditions.

**Figure S2:**
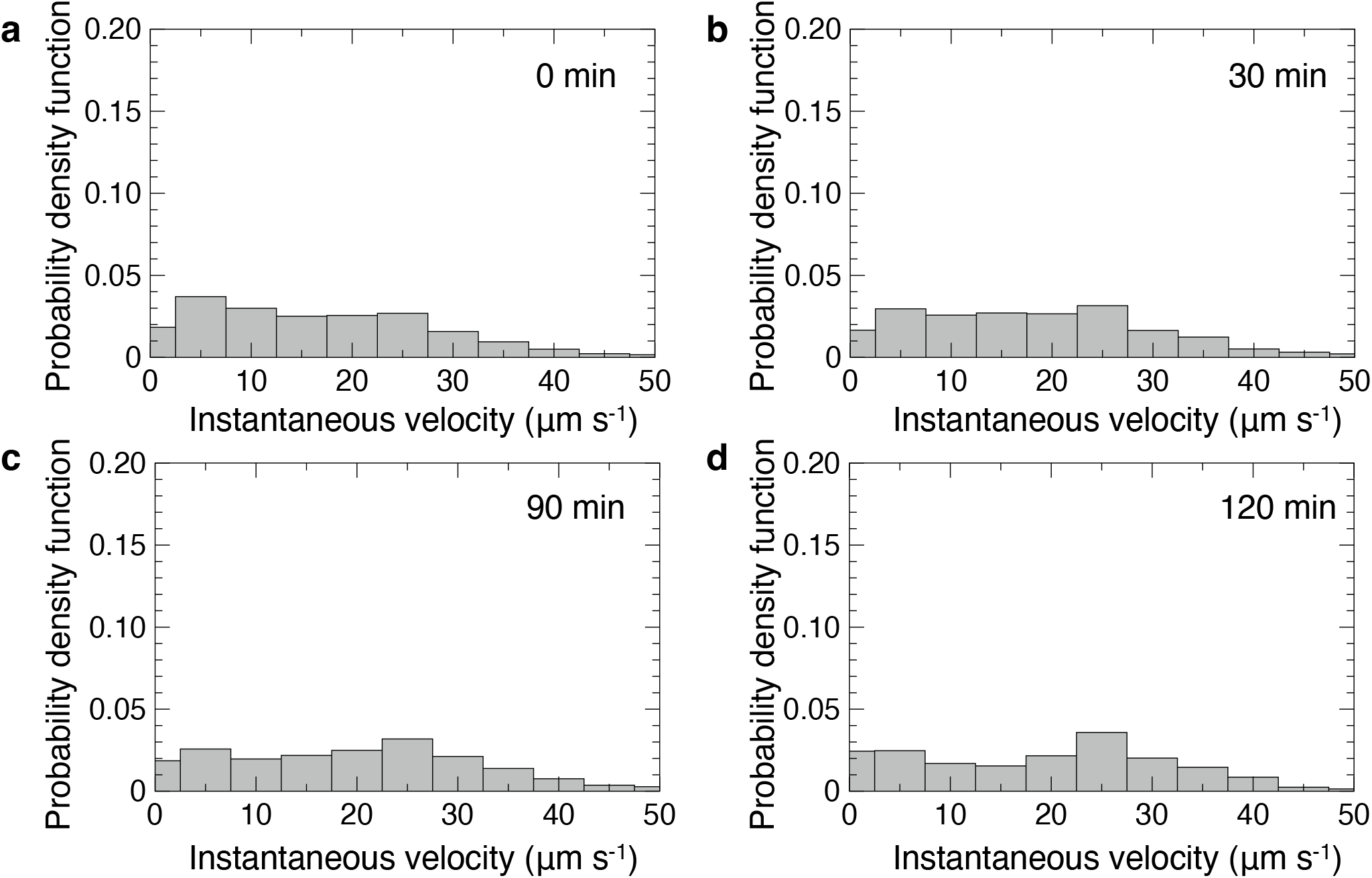
*E. coli* velocity in bulk BMB. (a–d) Probability distribution functions of instantaneous velocities for *E. coli* swimming in Berg’s Motility Buffer (BMB) at (a) 0 min, (b) 30 min, (c) 90 min, and (d) 120 min after loading. BMB contains no chemoattractant or nutrients, yet cells remain motile for at least 2 hours by using endogenous energy reserves.

### Note S4: Spreading rate experiment and image analysis

The population-scale spreading assays quantify how bacterial inocula expand through the transparent soils over time. We injected 2 *µ*L of a concentrated *E. coli* suspension (~ 1 × 10^9^ cells mL^−1^) into soil samples fully saturated with BMB. We introduced the bacteria directly at the center of the field of view and initiated imaging immediately using either a 2 × (N.A. 0.1) or 4 × (N.A. 0.2) objective. The optical configuration comprised a 153 *µ*m pinhole aperture and 488 nm excitation, capturing single optical sections ~ 100 *µ*m above the glass bottom. The image field of view was 8.8 × 8.8 mm for the 2× objective and 4.4 × 4.4 mm for the 4× objective. We recorded timelapse sequences spanning approximately 1 hour with 20 s frame intervals. For each frame, we calculated radially averaged fluorescence profiles centered at the inoculation site. At each timepoint, we generated 1 pixel-wide concentric annuli and recorded the mean pixel intensity within each annulus, yielding a profile *I*(*r*) as a function of radius *r*. We also determined a background intensity *I*_bg_ from regions well beyond the visible spreading front in the initial frame. A major analytical challenge is that the spreading dynamics are subdiffusive in the cryolite packings (see Figure S3), so extracting a physically meaningful diffusivity from population-scale profiles is not appropriate. Instead, we track the displacement of the leading edge of the population by defining a target intensity *I*_*c*_ = 1.05 *I*_bg_ and recording the radius *R*(*t*) where *I*(*r*) crosses this value.

**Figure S3:**
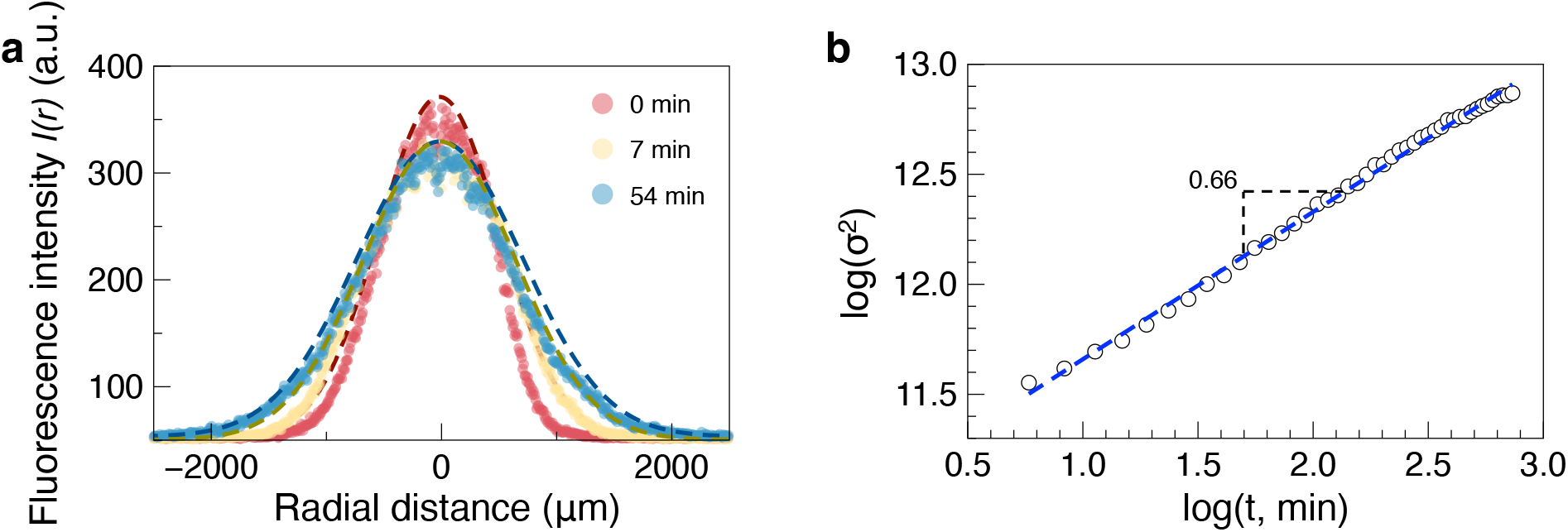
(a) Radial fluorescence profiles of a spreading *E. coli* inoculum in soil at representative timepoints, with Gaussian fits (colored lines). (b) Log-log plot of the squared profile width (variance) versus time, showing subdiffusive scaling (*σ*^2^ ∝ *t*^*α*^ with *α <* 1). The sublinear scaling precludes meaningful extraction of a diffusion coefficient at the population scale.

The trajectory *R*(*t*) of the leading edge typically shows an initial rapid quasi-linear expansion followed by a pronounced plateau. This behavior is predicted analytically for a spreading Gaussian from a standard standard solution to the diffusion equation: as material depletes near the origin, the front determined from a constant threshold must ultimately slow and plateau (Figure S4). This is shown as follows:

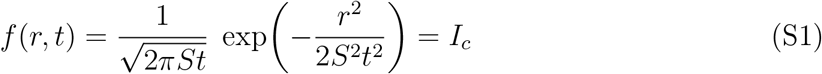

where *S* is the bacterial spreading rate. Solving for *r* = *R*(*t*):

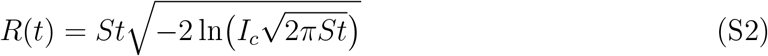

Showing how, for short times, the log term changes slowly such that *R*(*t*) ≈ *St*, i.e., an approximately linear front propagation at spreading rate *S*. We extract *S* by performing a piecewise linear fit on *R*(*t*), using findchangepts in MATLAB to identify the transition between the initial linear regime and the subsequent plateau.

**Figure S4:**
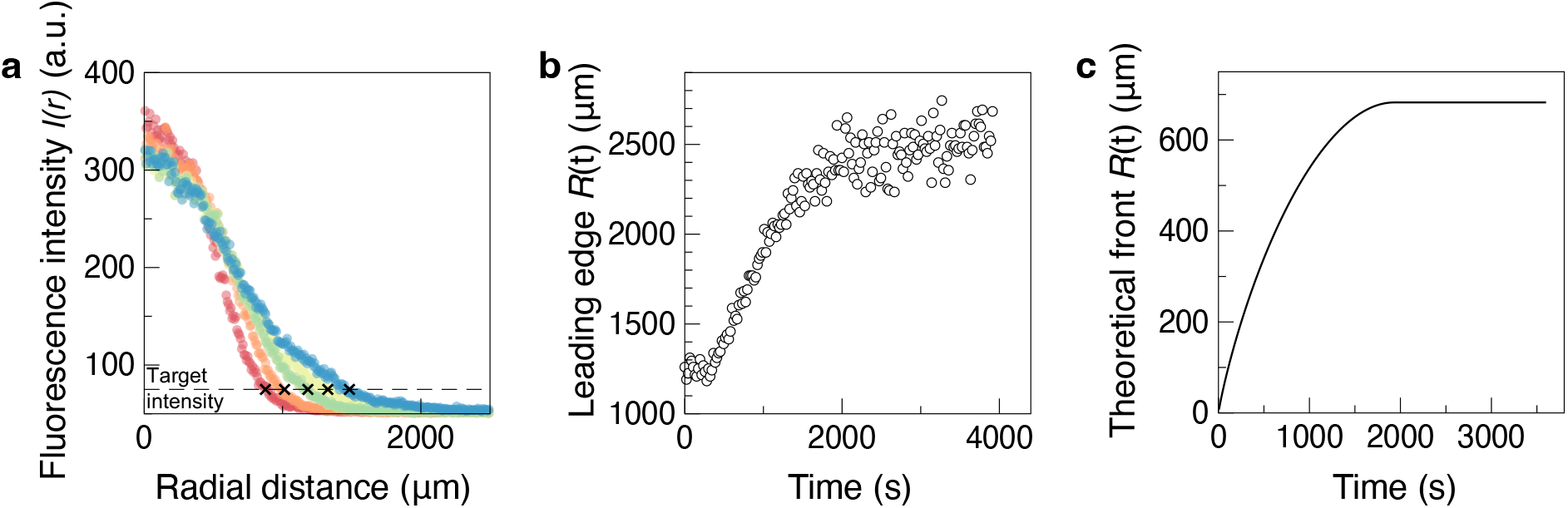
(a) Radial fluorescence profiles at successive timepoints, with the leading edge obtained from the target intensity crossing indicated by black cross symbols. (b) Time evolution of the leading edge position *R*(*t*) illustrating initial linear expansion and later plateau. (c) Analytical prediction of the leading edge position for a spreading Gaussian, demonstrating the expected plateau at long times due to mass conservation. The initial slope is used to define the population-scale spreading rate *S*.

### Note S5: Run vs trap detection algorithm

Quantitative analysis of pore scale bacterial motion required higher temporal resolution when recording fluorescence movies. We acquired timelapses using the resonant bidirectional scanning mode, with a 20× objective (N.A. 0.8), imaging with 0.86 × 0.86 *µ*m lateral resolution and at least 40 ms frame intervals. We tracked the position of individual bacteria using the TrackMate plugin in Fiji/ImageJ which extracts coordinates **x**(*t*) = [*x*(*t*), *y*(*t*)] and links them across successive frames from timelapse fluorescence microscopy movies. For each track, we applied a centered moving average filter with a fixed window of 5 timepoints to reduce noise before implementing a two-stage algorithm to classify bacterial motion into running and trapping states. For each timepoint *t*, we computed the displacement ratio *δ*(*t*) which is a scalar measure comparing the net displacement over a fixed interval to the total cumulative displacement over the same window:

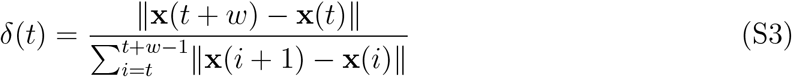

where **x** is the position vector of the moving bacterium and *w* is a fixed time window of 20 frames. The displacement ratio approaches 1 for directed motion and drops toward 0 for trapped, jitter-dominated motion. Any frame where *δ*(*t*) dropped below a fixed threshold *δ*_thresh_ was labeled as ‘trapped’. For frames not flagged as trapped, we applied a secondary check based on instantaneous kinematics. We computed the instantaneous velocity 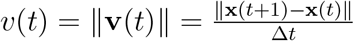 and the reorientation angle 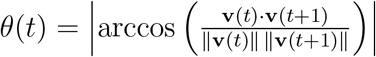. We classified a frame as ‘running’ only if *v*(*t*) *> v*_thresh_ and *θ*(*t*) *< θ*_thresh_. Frames were otherwise labeled as ‘trapped’.

We determined the input parameters of the classification algorithm, *w, δ*_thresh_, *v*_thresh_, *θ*_thresh_, and the smoothing window, through optimization of its precision and recall against manually labeled data. Precision is here defined as the fraction of predicted ‘running’ frames that are classified correctly, and recall is the fraction of manually labeled ‘running’ frames that are successfully identified by the algorithm. We independently annotated a subset of tracks frame-by-frame as ‘running’ or ‘trapped’ based on visual inspection and applied the classification algorithm while systematically varying each of the parameters. We then computed the F1 score to quantify the tradeoff between classification precision and recall:

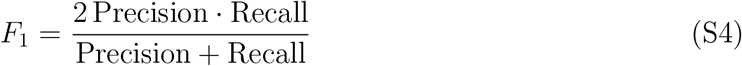

We selected the following parameters that maximize F1 score, as shown in Figure S5: *w* = 20 frames; *δ*_thresh_ = 0.6; *v*_thresh_ = 6.5 *µ*m s^−1^; *θ*_thresh_ = *π/*6; Position smoothing window = 5 frames.

**Figure S5:**
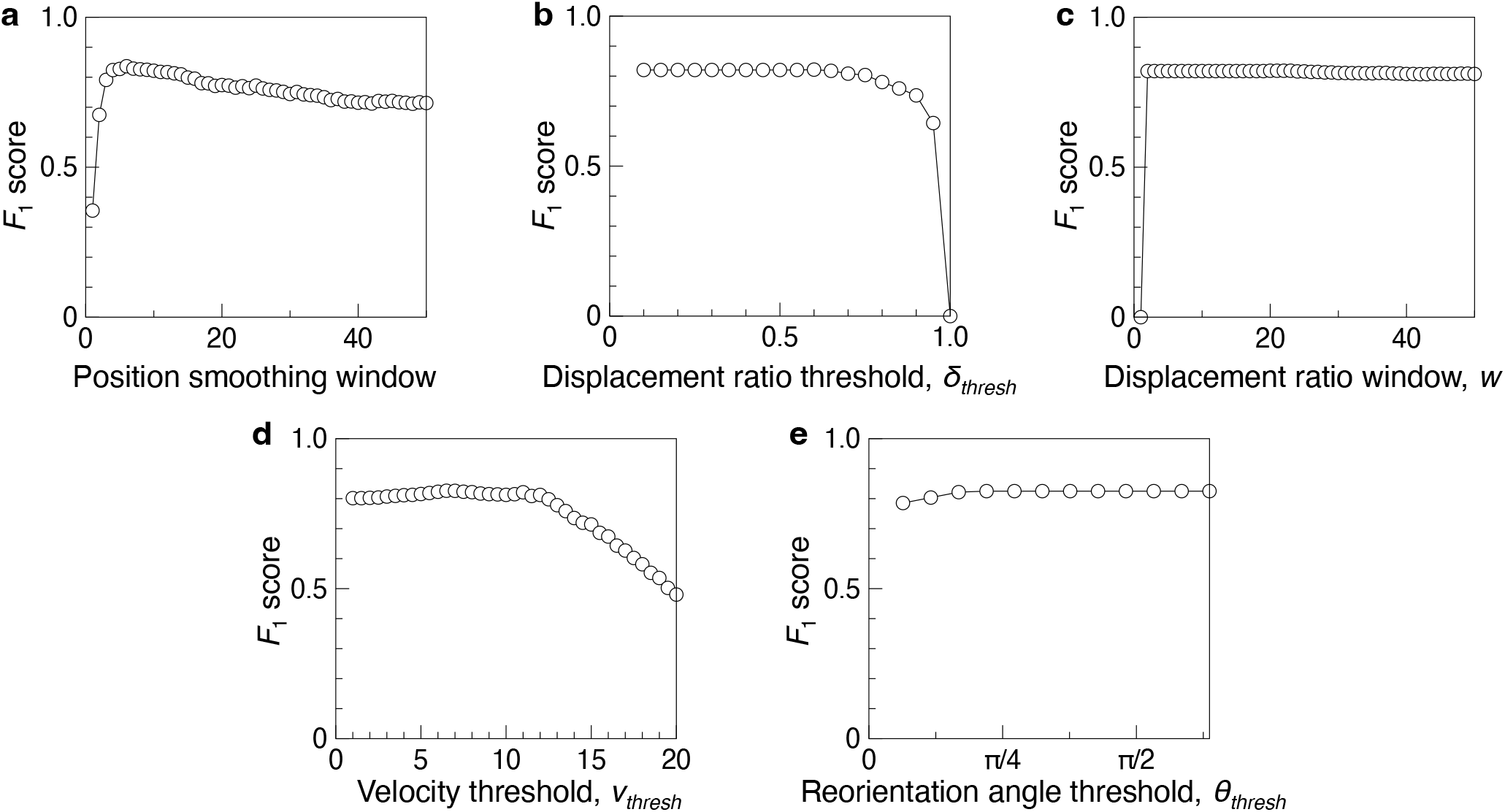
Parameter sensitivity of the *F*_1_ score evaluated against manual ground-truth labeling. (a) Position smoothing window, i.e., number of frames for centered moving average. (b) Displacement ratio threshold for distinguishing directed motion from trapped states. (c) Displacement ratio window size, i.e., number of frames used for displacement ratio calculation. (d) Velocity and (e) reorientation angle thresholds used to refine classification.

**Figure S6:**
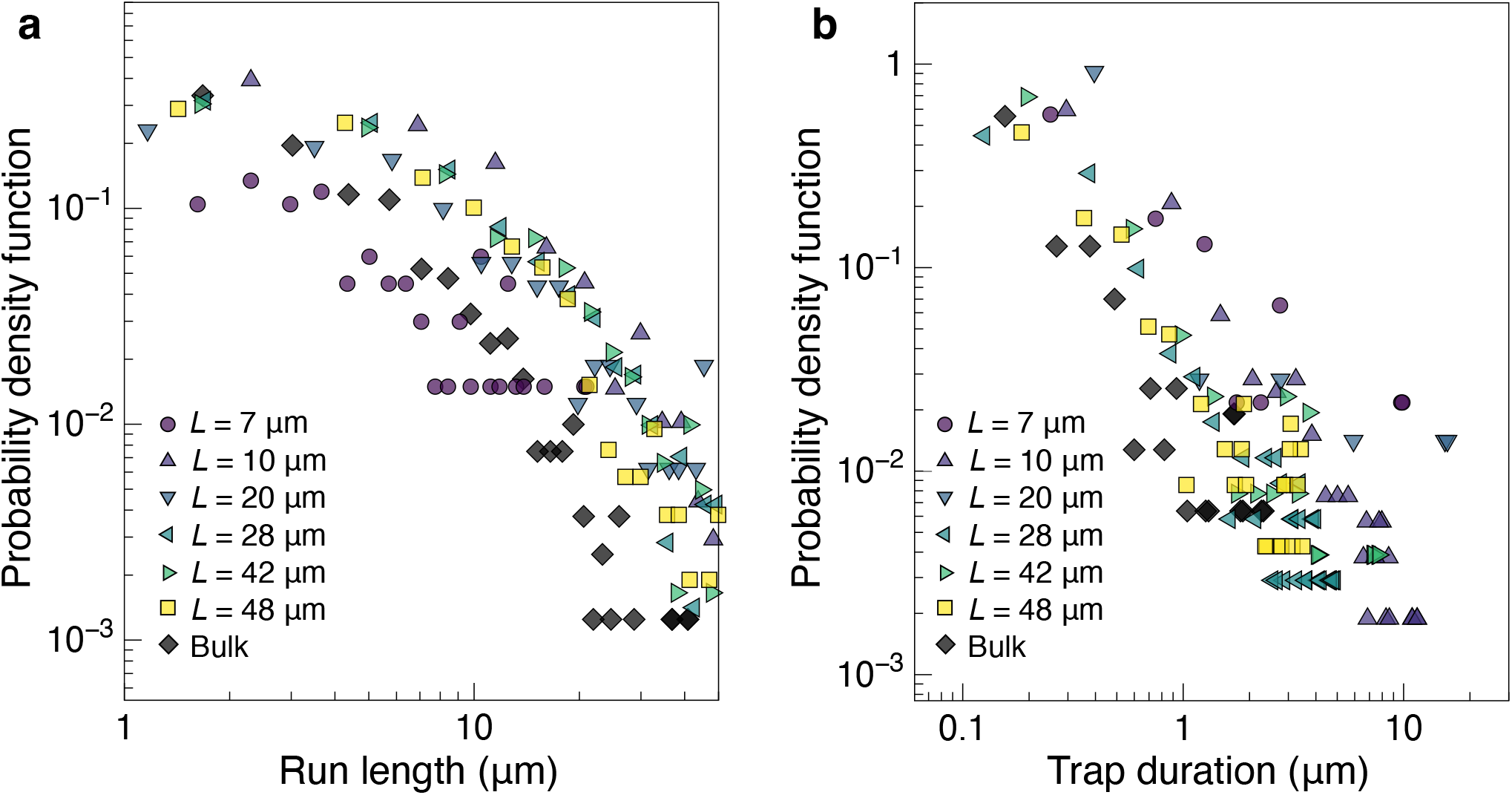
Probability density functions of (a) run length and (b) trap duration for each soil texture and bulk BMB media. Measurements of run length are limited to segments that are bounded by two trapped states. Similarly, measurements of trap duration are limited to periods bounded by running states.

**Figure S7:**
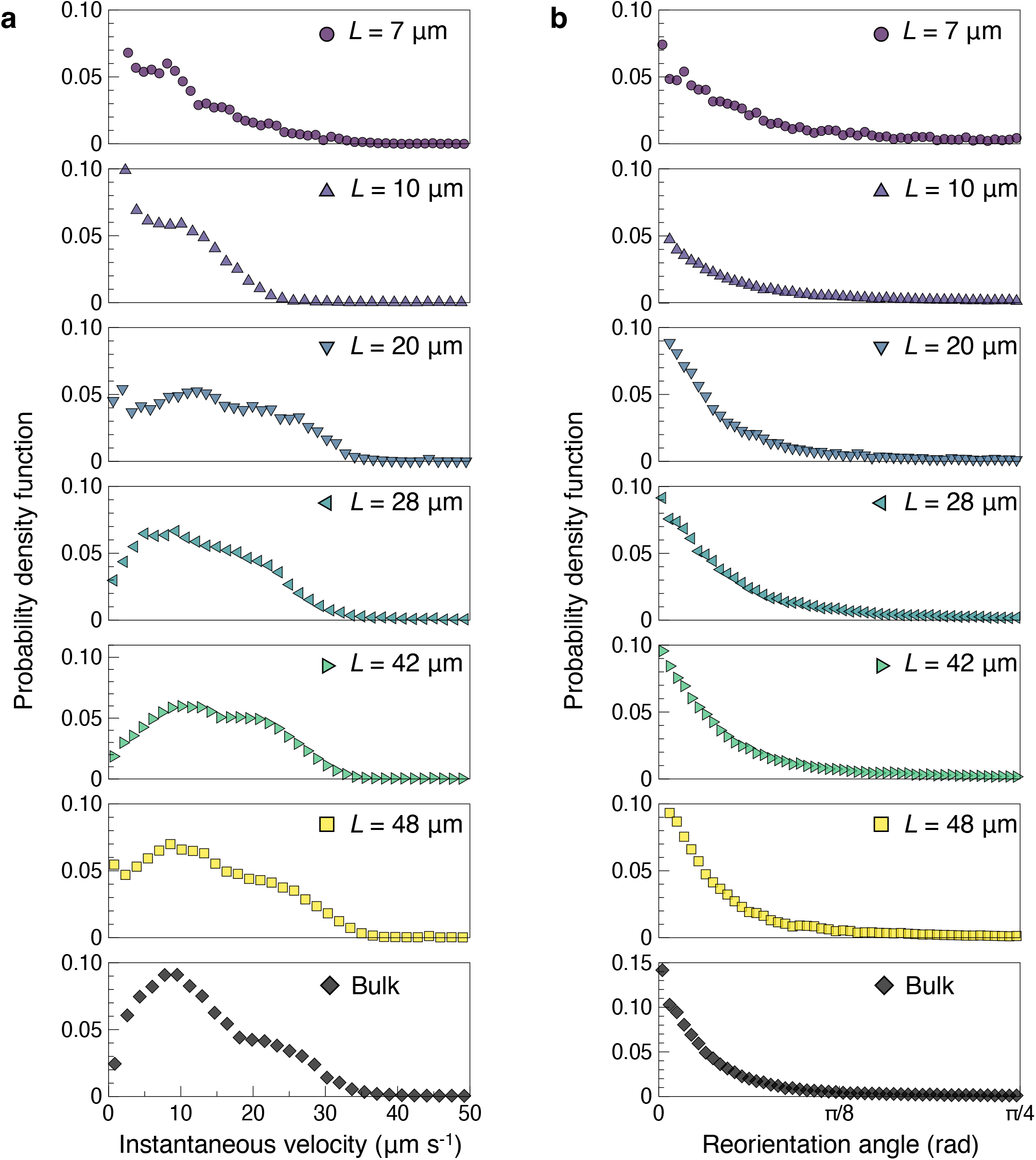
Probability density function of (a) instantaneous velocity and (b) reorientation angle for each soil texture and bulk BMB media.

### Note S6: Mean squared displacement analysis

We computed the time-averaged mean squared displacement (MSD) curves from individual trajectories which is calculated as follows:

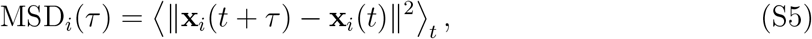

where **x**_*i*_(*t*) is the position vector of bacterium *i* at time *t*, and *τ* is the lag time. The ensemble MSDs plotted in Figure 3 emerge from averaging individual MSDs:

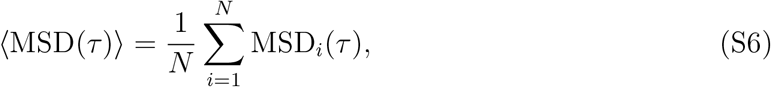

with standard error of the mean computed across tracks at each lag time. We then extract the crossover point between motility regimes via segmented linear fitting in log-log space. Two separate fits were applied to the MSD curve over the ballistic regime typically between *τ*_1_ ≈ 0.1 s and *τ*_2_ ≈ 0.8 s, and the diffusive regime typically between *τ*_3_ ≈ 1 s and *τ*_4_ ≈ 3 s:

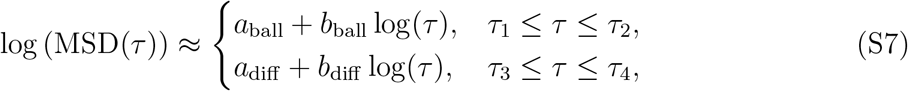

where the subscripts ‘ball’ and ‘diff’ reflect the ballistic and diffusive regimes, respectively. We obtain the crossover lengthscale *λ*_*c*_ after extracting the crossover lag time *τ*_*c*_:

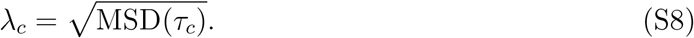

Finally, we estimated the effective diffusivity *D*_eff_ by fitting a linear model to the MSD curve over the diffusive range:

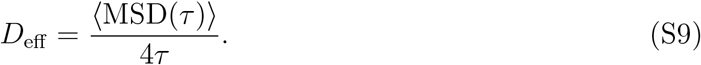

**Figure S8:**
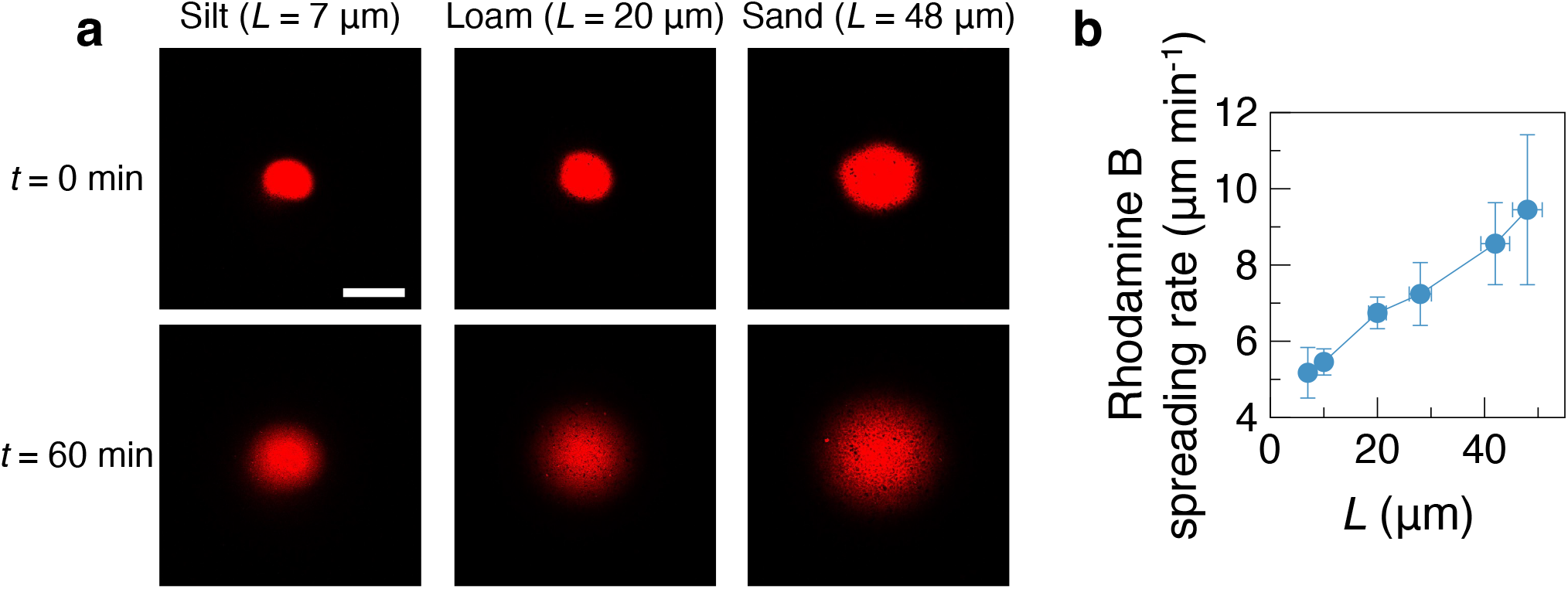
Rhodamine B spreading rates in transparent cryolite soils. (a) Representative confocal micrographs of Rhodamine B droplets (red) injected into cryolite packings with characteristic pore lengths *L* = 7 *µ*m (silt), *L* = 20 *µ*m (loam), *L* = 48 *µ*m (sand). Top row: immediately after injection; bottom row: after 60 min of spreading. Scale bar: 1 mm. (b) Measured spreading rates of Rhodamine B versus characteristic pore size *L* for three biological replicates per texture. Data points and vertical error bars indicate mean ± standard deviation. Horizontal error bars represent the 95% confidence interval from exponential fits to the pore chord length distributions used to extract *L*.

**Figure S9:**
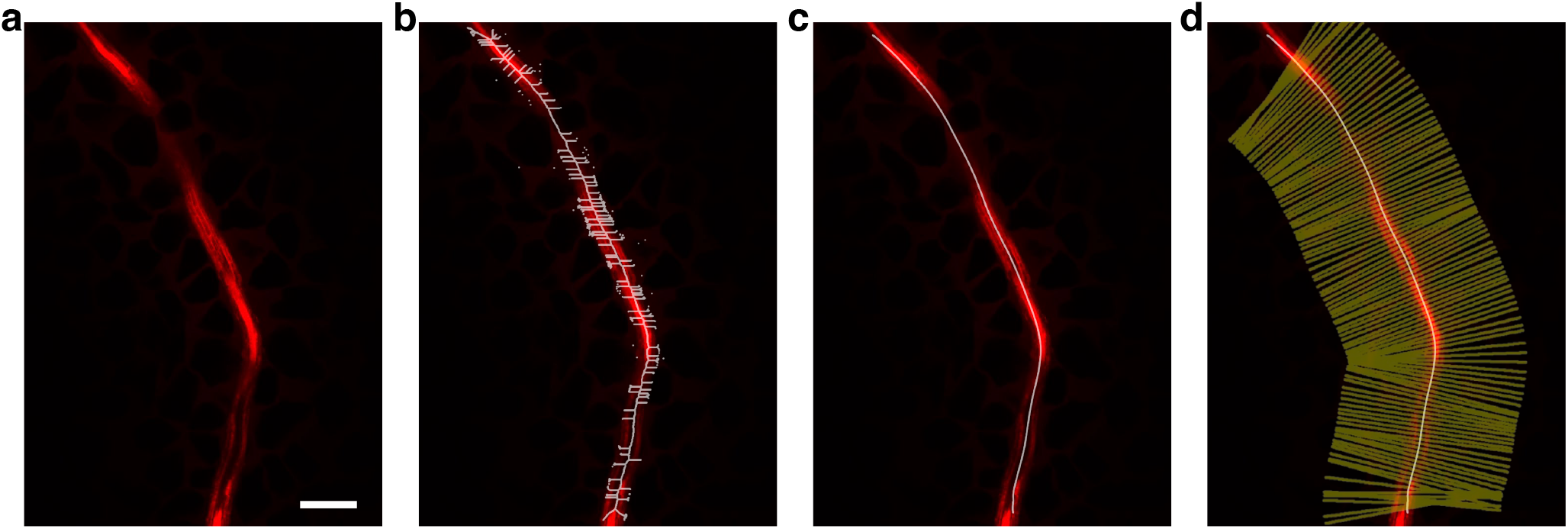
Workflow for extracting fluorescence intensity profiles around curvilinear individual root. (a) Confocal fluorescence image of a root (red) embedded in a transparent cryolite matrix. (b) Skeletonization of the root region yields a pixel-wide medial axis and small spurious branches. (c) Centerline is extracted as the longest branch of the skeleton and smoothed with a moving average filter (100 *µ*m window size). (d) Sampling grid is constructed from outward normals (yellow) to the segmented root boundary, oriented by the smoothed centerline. Distance *d* is measured along each normal starting at the root surface and extending into the soil. Scale bar: 200 *µ*m.

**Figure S10:**
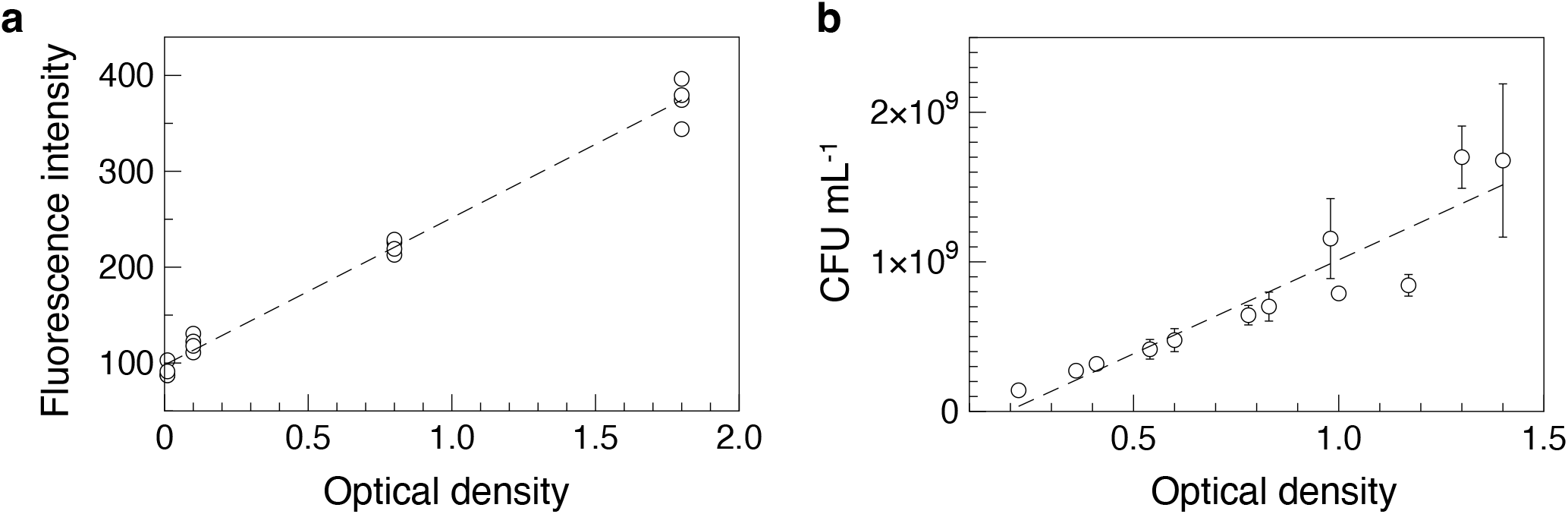
Calibration of fluorescence intensity to cell concentration. (a) Fluorescence intensity vs optical density (OD); open circles are individual measurements and the dashed line is the linear fit, *I* = 154 *OD* + 98. (b) Colony forming units per milliliter (CFU mL^−1^) vs OD; open circles show measurements (error bars indicate standard deviation across replicates, *n* = 3), and the dashed line is the linear fit, CFU = 1.26 × 10^9^ *OD* − 2.42 × 10^8^. Combining the two fits gives a direct fluorescence to concentration conversion, CFU = 8.2 × 10^6^*I* − 1 × 10^8^.

